# Cis-regulatory divergence and misexpression of spermatogenesis genes underlie hybrid male sterility in the *Drosophila bipectinata* species complex

**DOI:** 10.1101/2025.09.18.677105

**Authors:** M Manjunath, C S Damini, V Shakunthala

**Affiliations:** Current address: Research Scholar, Chronobiology lab, Department of Studies in Zoology, University of Mysore, Manasagangothri, Mysuru 570006, Karnataka, India; Current address: Professor, Department of Studies in Zoology, University of Mysore, Manasagangothri, Mysuru 570006, Karnataka, India

**Keywords:** Hybrid male sterility, Spermatogenesis, Gene expression, Regulatory divergence, Cis-regulatory motifs

## Abstract

Hybrid male sterility (HMS) is one of the most common and earliest forms of postzygotic reproductive isolation in *Drosophila*, often arising from defects in spermatogenesis linked to regulatory divergence of rapidly evolving male-biased genes. The *Drosophila bipectinata* species complex (*D. bipectinata, D. malerkotliana, D. parabipectinata, and D. pseudoananassae*) provides a valuable system to study the speciation paradigm, since, all the twelve F₁ hybrid males are sterile, the precise stage of spermatogenic failure differs among crosses. Here, we integrate cytological, transcriptional, and regulatory sequence analyses to examine the basis of sterility in this sub-complex. Cytological assays revealed that hybrid testes exhibit developmental arrest at different spermatogenesis stages, covering from reduced primary spermatocytes to absence of individualized sperm. Quantitative RT-PCR of seven spermatogenesis genes—*aly*, *bam*, *sa*, *dj*, *topi*, *can*, and *Mst98Ca*—showed significant downregulation among hybrid males relative to parental species, linking transcriptional misregulation with observed phenotypic defects in spermatogenesis. To test whether regulatory divergence underlies this misexpression, we analyzed ∼1–2 kb promoter regions of five genes (*aly*, *bam*, *sa*, *dj*, and *topi*). Motif scans using FIMO identified several lineage-specific turnover of transcription factor binding sites, also PWM-delta analysis revealed substantial interspecific differences in predicted binding strength for many key testis transcription factors. These results demonstrate that HMS in the *D*. *bipectinata* complex is consistently associated with spermatogenic arrest and misregulation of genes essential for germline proliferation, meiotic progression, and spermiogenesis. The concordance between expression changes and promoter divergence supports a role for cis-regulatory evolution in hybrid dysfunction, while also leaving open the contribution of trans-acting factors. This study situates the *D*. *bipectinata* complex within the broader framework of *Drosophila* hybrid sterility, extending and highlighting the evolutionary sensitivity of spermatogenesis and regulatory divergence in speciation.

## Introduction

Reproductive isolation is a central mechanism in speciation, and hybrid male sterility (HMS) is one of the most common and informative forms of postzygotic isolation in most of the sexually reproducing animals with heterogametic males including *Drosophila* (Haldane, 1922; Presgraves, 2010; Presgraves and Meiklejohn, 2021). HMS often arises from defects in the spermatogenesis pathway, in which developmental stages getting arrested is closely associated with widespread misregulation of spermatogenesis genes in hybrid testes (Michalak and Noor, 2003; Moehring et al., 2007; Civetta, 2016). Such sterility reflects the combined action of Dobzhansky–Muller incompatibilities, rapid evolution of male-biased genes, and divergence in cis-and trans-regulatory elements (Wu and Davis, 1993; Masly and Presgraves, 2007).

Genes expressed in the male reproductive tract are well known to evolve rapidly. From early proteomic surveys which revealed higher interspecific divergence in the reproductive tract proteins compared to other tissues (Coulthart and Singh, 1988; Thomas and Singh, 1992; Civetta and Singh, 1995), to comparative genomics of 12 *Drosophila* species revealed that male-biased genes not only exhibit elevated sequence divergence but also display higher rates of ortholog loss across lineages, particularly for genes critical to gametogenesis (Haerty et al., 2007). Expression studies have further added this narrative by showing that male-biased genes diverge more strongly in transcriptional profiles between species than female-biased or unbiased genes (Meiklejohn et al., 2003; Ranz et al., 2003; Zhang et al., 2007). These findings underpin the “faster-male” hypothesis, in which the rapid evolution of male-biased genes and their regulatory elements predisposes them to mismatches in hybrids (Assis et al., 2012; Grath and Parsch, 2012). As a consequence, HMS has been a recurrent outcome of interspecific crosses in *Drosophila* and a central focus for testing evolutionary theories of speciation. Cytological studies have long reported profound disruptions in spermatogenesis in sterile hybrids. Early work by Dobzhansky (1934) documented hybrid testes containing spermatocytes that entered meiosis but failed to complete differentiation into motile sperm. Similar observations were extended in the *Drosophila simulans* clade, where sterile hybrid males between *D. simulans*, *D. mauritiana*, and *D. sechellia* exhibited defective onion cell formation, disorganized cysts, and arrested spermatid differentiation (Lachaise et al., 1986; Wu et al., 1992; Kulathinal and Singh, 1998). Strikingly, sterility-associated spermatogenic defects are not unique to *Drosophila*: other individuals such as sterile hybrid Xenopus males also have shown massive reductions in sperm counts and abnormal spermatid morphology (Malone et al., 2007). These consistent hybrid male sterility paradigm among the different taxa suggests that gametogenesis is particularly a fragile process during hybridization. In *D. melanogaster*, spermatogenesis proceeds through a complex sequence of mitotic proliferation, meiotic divisions, and spermiogenesis, with distinct sets of genes regulating each stages particularly (Fuller, 1993; Fuller, 1998; White-Cooper et al., 2009). Genes such as *bam* and *bgcn* regulate the switch from mitosis to meiosis transition, aly and sa are key members of the meiotic arrest gene family required for progression into meiosis, and *dj*, *topi*, *can*, and *Mst98Ca* control later stages of spermatid differentiation and individualisation (Fuller, 1998; Jiang and White-Cooper, 2003; Santel et al., 1998). Comparative studies in the *D. simulans* clade have revealed ubiquitous misregulation of these candidate spermatogenesis genes in sterile hybrids, with a strong bias toward downregulation of post-meiotic spermiogenesis genes (Michalak and Noor, 2003; Moehring et al., 2007; Catron and Noor, 2008; Michalak and Ma, 2008; Sundararajan and Civetta, 2011). Also, early-acting regulators such as bam and sa have also been shown to be being misregulated in testes-specific analyses (Sundararajan and Civetta, 2011), emphasizing the importance of tissue-specific assays. These results together highlight both sterility-specific misregulation and faster-male regulatory divergence as overlapping processes shaping expression patterns in hybrids.

Despite the detailed work in the *melanogaster*–*simulans* clade, other *Drosophila* species complexes and lineages remain comparatively underexplored at the intersection of cytological, transcriptional, and regulatory analyses. The *Drosophila bipectinata* species complex, which includes *D. bipectinata*, *Drosophila malerkotliana*, *Drosophila parabipectinata*, and *Drosophila pseudoananassae*, is particularly intriguing because all F₁ male hybrids are sterile, although the precise developmental stage of arrest varies depending on the parental species combination (Mishra and Singh, 2006). These patterns echo the variation seen in the *simulans* clade and suggest that multiple incompatibilities act at different points of spermatogenesis. Yet the molecular correlates of these sterility phenotypes have not been systematically investigated in this group. To address this, we combined cytological observations, expression assays, and promoter sequence analyses in the *bipectinata* complex. Transcript levels were quantified by qRT-PCR for seven spermatogenesis genes spanning different developmental stages (*aly, bam, sa, dj, topi, can,* and *Mst98Ca*), while promoter motif and PWM-delta analyses were conducted for those genes with validated upstream sequences (*aly, bam, sa, dj,* and *topi*). To ensure statistical robustness, each comparison of testes RNA between sterile hybrids and fertile parental species was conducted with five biological replicates, and expression levels were normalized using a single housekeeping gene. Across all the assays, a general trend of downregulation was observed for most of the genes examined in several samples of sterile hybrids relative to their parental species. This reduction was statistically significant for genes involved in germline proliferation, meiotic control, and spermatid differentiation. and importantly, the downregulation was detected in the RNA extracted from testes, but not from whole-fly or ovarian RNA samples. These findings suggest that the observed expression changes may arise from male-specific divergence in regulatory elements or may reflect downstream consequences of subtle developmental defects in sterile males. These complementary approaches link phenotypic arrest in spermatogenesis with both transcriptional changes and underlying regulatory sequence divergence, providing a basis for evaluating whether misexpression in sterile hybrids reflects sterility itself, faster-male regulatory evolution, or both.

## Materials and methods

### *Drosophila* stocks and Hybridization regime

The stocks employed during present study were *D. bipectinata* Pune, *D. parabipectinata* Mysore, *D. malerkotliana malerkotliana* Raichur (all from India) and *D. pseudoananassae* nigrens Brunei Island, Brunei (procured from the BHU, Varanasi, India). All the stocks were maintained in the laboratory on simple yeast agar medium at approximately 240 C with a relative humidity of 70% in a 12 h dark; 12 h light cycle, by transferring approximately 40 flies (males and females in equal number) to fresh culture bottles in every generation. To obtain hybrid males we intercrossed virgin females of one species with virgin males of heterospecific sister species (7 days aged virgin flies as parents) in no choice mating combination, while the combination of both conspecific and heterospecific males with a single female failed to produce any hybrids as the female always chose conspecific males (Supporting table 3). When progeny emerged, males and females were sorted and kept in separate vials. To study the development of spermatogenesis 50 matured male hybrids (7-10 days old) were randomly selected for observation.

### Light and Phase contrast microscopy

To study the developmental stages of spermatids and to observe the motility of sperm in parental and hybrid male flies, testes along with seminal vesicles separated from the accessory gland and external genitalia were mounted and gently squashed under a cover slip in the PBS drop following the method of Kemphues et al., (1980) and were observed under light microscope. Ten males from each parental *Drosophila* species and a minimum of 20 males from each hybrid were analyze*D.* Dissections were performed in a droplet of 1× PBS, and any abnormalities in testes morphology were recorde*D.* The seminal vesicle was punctured to release sperm, and sperm motility was evaluated using a phase-contrast microscope and the images were taken at 40-100 x magnification.

### Selection of genes

Candidate genes for gene expression assays were chosen from a curated set of genes previously evaluated in *Drosophila* sterile hybrids. Initially, we prioritized genes analyzed in qRT-PCR assays, focusing on those exhibiting misexpression in sterile male hybrids. Additionally, we selected genes identified as underexpressed in interspecies sterile hybrids of the *simulans* clade (Moehring et al., 2007). From this pool, we restricted our targets to genes functionally annotated for spermatogenesis or spermatid development in AmiGO (http://amigo.geneontology.org/). To ensure specificity, we excluded genes lacking orthologs in the *D. bipectinata* subcomplex, as well as those with multiple orthologs, thereby increasing confidence in amplifying a single target gene. Orthologs of *D. bipectinata* genes expressed during spermatogenesis were identified using OrthoDB Orthologs within FlyBase (http://flybase.org/).

### Gene expression analysis

Virgin males of pure and hybrids of *D. bipectinata* species complex, were aged to sexual maturity (as previously described), and their testes, excluding accessory glands but including seminal vesicles, were dissected and rinsed in 1× PBS. Five biological replicates were prepared for each parental species and hybrids, with each replicate comprising ten (Pure and hybrid group) testes. Total RNA was extracted through phenol-chloroform method, and quantified via a nanophotometer. Complementary DNA (cDNA) was synthesized from standardized amounts of total RNA using the iScript cDNA Synthesis Kit.

Primers were designed with Primer-Blast (https://www.ncbi.nlm.nih.gov/tools/primer-blast/) to generate amplicons ranging from 75 to 120 base pairs. Whenever feasible, intron-spanning primers were employed to facilitate detection of genomic DNA contamination (Table S2). Testicular expression of selected candidate genes was initially validated using standard PCR, performed with a 5-minute initial denaturation at 95°C, followed by 35 cycles of 45 seconds at 95°C, 30 seconds at 60°C, and 45 seconds at 72°C, concluding with a 15-minute extension at 72°C. PCR products were visualized on a 2% agarose gel to confirm expression. Subsequently, gene expression was quantified via qRT-PCR using the IQ SYBR Green Quantitative Real-Time PCR Kit (Bio-Rad). One reference gene, RpL32 were used to normalize qRT-PCR results for each target gene. The qRT-PCR cycling conditions were uniform across all genes: an initial 5-minute denaturation at 95°C, followed by 35 cycles of 45 seconds at 95°C, 30 seconds at 60°C, and 45 seconds at 72°C. Gene expression levels were calculated using the ΔCT method, defined as the CT of the reference gene (RpL32) minus the CT of the target gene. All the parental species served as independent calibrators to compute ΔΔCT, determined as the difference between the ΔCT of each sample and the average parental ΔCT.

### DNA Extraction and Gene Sequencing

Genomic DNA was extracted from pools of ∼50 adult flies per strain of the *Drosophila bipectinata* species complex using the DNeasy Blood and Tissue Kit (QIAGEN), following the manufacturer’s instructions. Primers were designed for six spermatogenesis genes—*always early (aly)*, *bag of marbles (bam)*, *spermatocyte arrest (sa)*, *don juan (dj)*, and *matotopetli (topi)*—based on reference sequences from *D. bipectinata* (Supporting Table S1). The complete coding regions of these genes were amplified by polymerase chain reaction (PCR), and the resulting amplicons were purified and sequenced using Sanger sequencing to confirm gene identity and integrity.

### Sequence acquisition and promoter region assembly

For promoter motif analyses, ∼1–2 kb upstream sequences were obtained for each gene from all four species. Sequences were derived from de novo Sanger sequencing (this study) and from publicly available NCBI/FlyBase assemblies (NCBI Resource Coordinators, 2018; FlyBase Consortium, 2003) when validated reference sequences were not available. Contigs of multiple Sanger sequences were assembled using CAP3 (Huang & Madan, 1999), and sequences were manually curated to ensure correct open reading frame orientation.All the sequences are available at 10.6084/m9.figshare.30102763.

### Motif discovery and presence/absence analyses

Promoter regions were scanned for transcription factor binding sites (TFBS) using FIMO (Find Individual Motif Occurrences) in the MEME Suite v5.5.8 (Grant et al., 2011; Bailey et al., 2015). Position weight matrices (PWMs) were obtained from the JASPAR 2024 (Castro-Mondragon et al., 2022) insects database. Default background models were used for the analyses, and motifs with q ≤ 0.05 was considered significant. Presence/absence of motifs across species was summarized in a binary motif-by-species matrix. Heatmaps were generated in Python (matplotlib/seaborn) (Hunter, 2007; Waskom, 2021), the complete list of FIMO hits is provided in Supporting Table S4.

### PWM-delta analysis

To quantify interspecific differences in predicted motif strength, PWM-delta analysis was performe*D.* For each motif × gene combination, the highest FIMO PWM score per species was extracte*D.* The difference between maximum and minimum scores across species (Δ) was used as a metric of regulatory divergence. Motifs with Δ > 5 were considered highly divergent. Barplots were generated for the top five divergent motifs (Fig 8), and the full PWM-delta dataset is available in Supporting Table S5.

### Statistical analysis and visualization

All statistical analyses were conducted using ANOVA and nonparametric models in Graph-pad PRISM (GraphPad Software, 2021) and Python’s SciPy package (Virtanen et al., 2020). Figs were generated using matplotlib (Python v3.11) and R ggplot2 (Wickham, 2016). Final Fig panels were assembled in Adobe Illustrator (Adobe Systems Inc., 2022)..

## Results

### Inter-species hybrid male sterility

Interspecific crosses within the *Drosophila bipectinata* sister species complex results in complete sterility, as hybrid males are incapable of producing viable offspring, whether mated with hybrid females or backcrossed with females of the parental species. We conducted the F1 hybrid matings and assayed the sperm storage organ (spermatheca) of the F1 female hybrids for the presence of sperms within one hour after copulation. But none of the female hybrids showed any sperms inside the spermatheca and seminal receptacle, which specifies that hybrid males have failed to transfer sperms or if there is a possibility of faster sperm spillage by the hybrid females within 30 minutes after copulation. Hence in order to check the motility of sperms in hybrid males testes along with seminal vesicles were observed through phase contrast microscopy, motile sperms were seen in the 6 days old testes of pure species respectively(Fig 1), but all the twelve different hybrid males failed to produce motile sperms. F1 hybrid males of all the heterospecific parental matings except those of the hybrid males having *D.pseudoannanassae* as female parent, failed to produce any individualised motile sperm(Fig 2). In the hybrids among which *D.pseudoannanassae* served as maternal parent, testis showed no spermatids in which spermatogenesis development has arrested in its preliminary stages itself(Fig 3).

**Fig 1:**
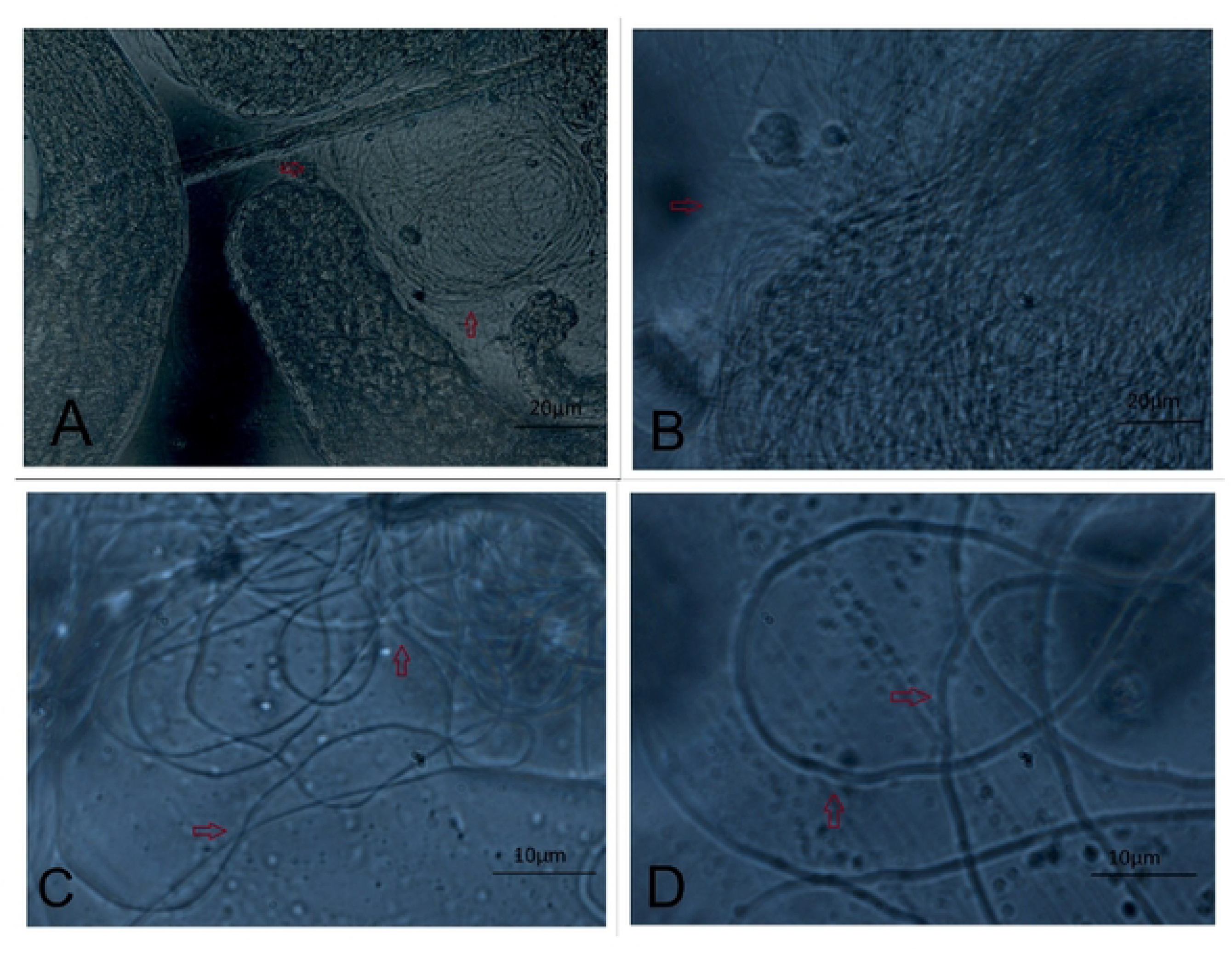
Phase contrast view of testes squash from 4-day old parental males showing individualised sperms (A) *D.bipectinata* (B) *D. malerkotliana* (C) *D.parabipectinata* (D)*D. pseudoananassae.* arrows indicate individualised sperms around the testes. (Magnification:40-100X / Scale bar:10 - 20μm).

**Fig 2:**
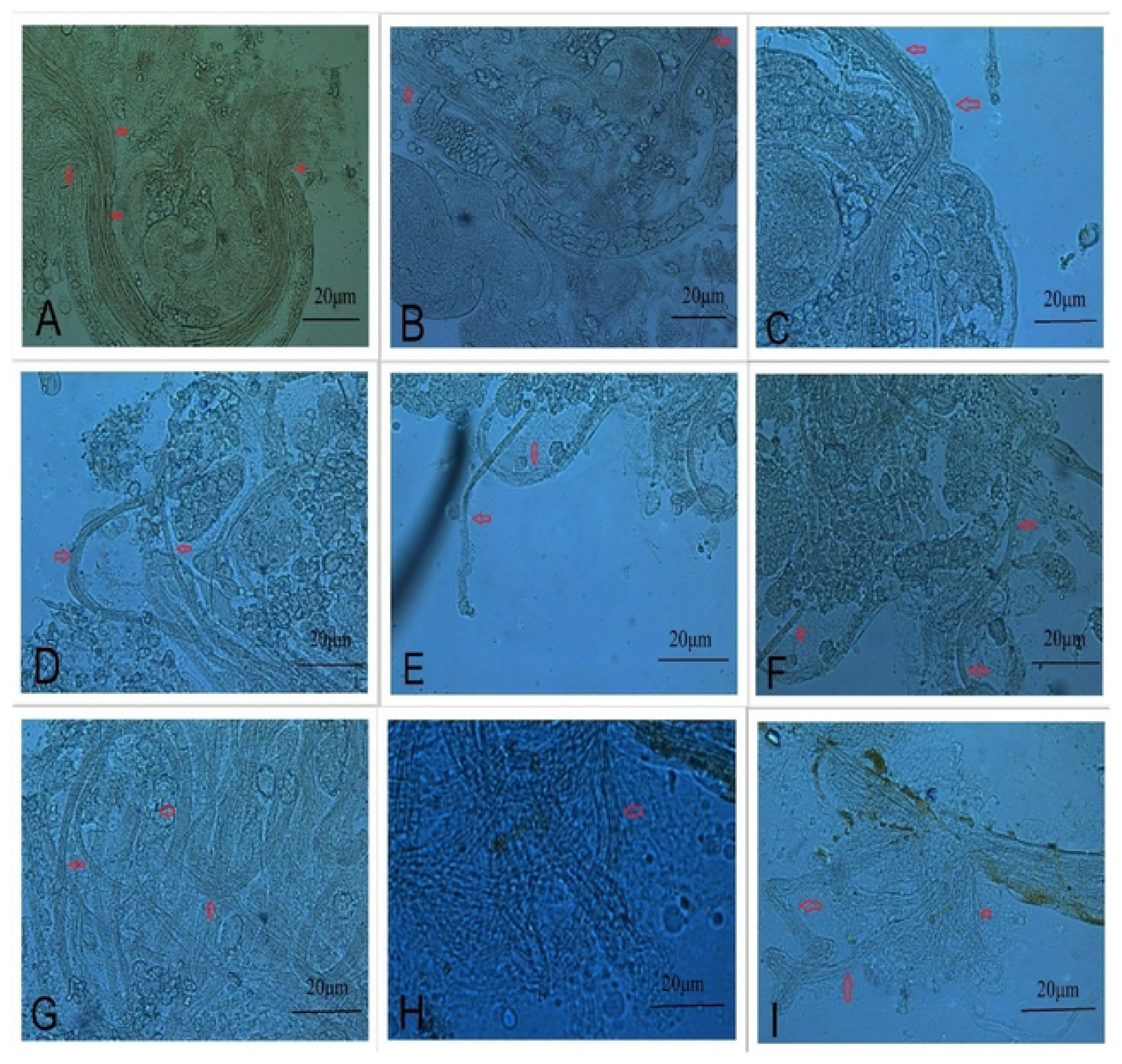
Phase contrast view of mid testes squash of post-meiotically defective hybrids showing unindividualised spermatids (A) *D. bipectinata♂ X D. malerkotliana♀* (B*) D.bipectinata♂ X D.parabipectinata♀ (*C*) D. malerkotliana♂ X D.bipectinatata♀ (*D*) D. malerkotliana♂ X D.parabipectinata♀ (*E*) D. pseudoananassae♂ X D.bipectinatata♀ (*F*) D. pseudoananassae♂ X D. malerkotliana♀ (*G*) D.parabipectinata♂ X D.bipectinatata♀ (*H*) D.parabipectinata♂ X D. malerkotliana♀ (*I*) D. pseudoananassae♂ X D. malerkotliana♀. (Magnification:40X / Scale bar:* 20μm).

**Fig 3:**
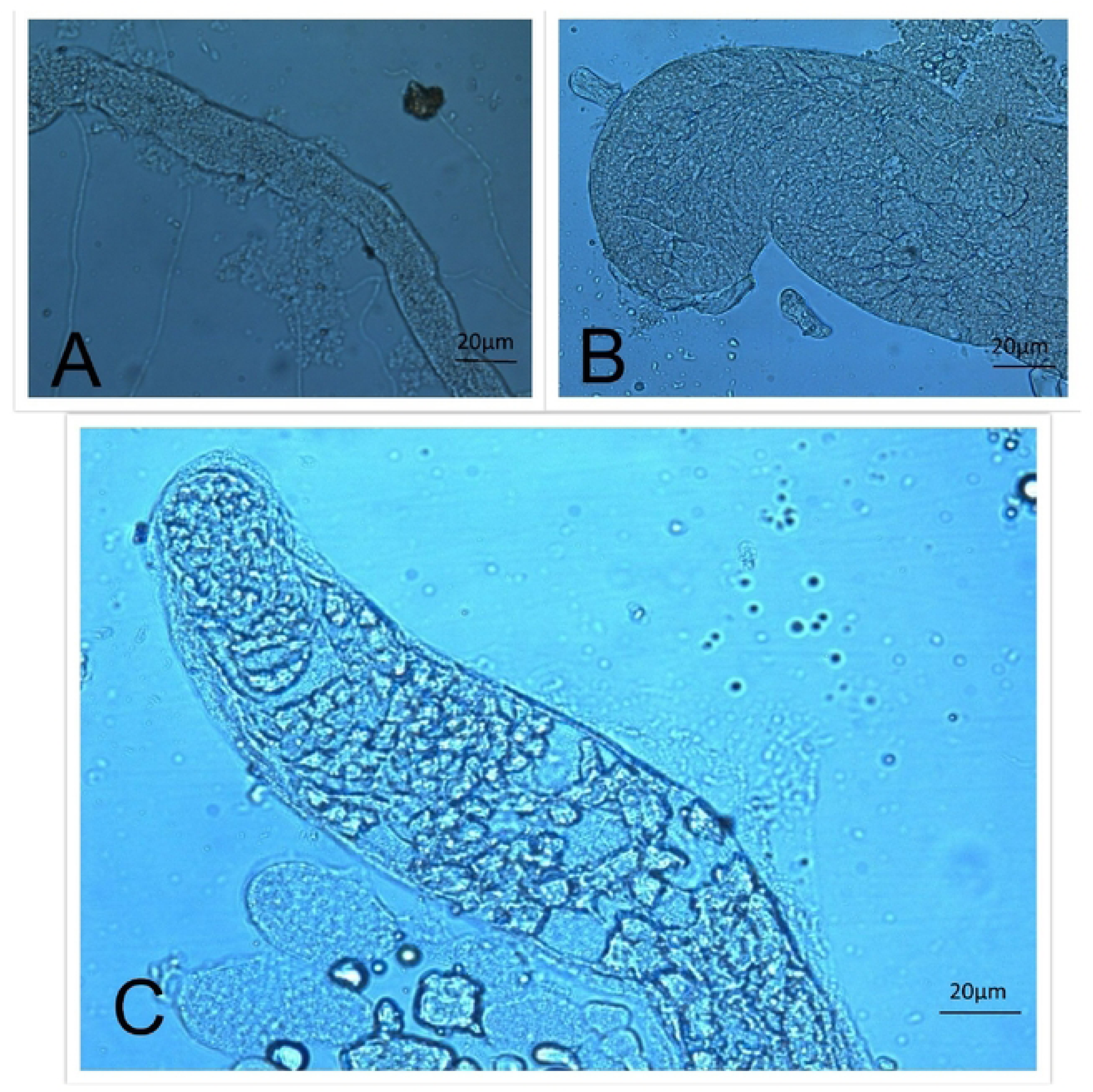
Aspermic testes which has arrested in the pre-meiotic stages where the germ cells have failed to differentiate into primary spermatocytes among the hybrids (A) *D. bipectinata♂ X D. pseudoananassae♀* (B) *D. malerkotliana♂ X D. pseudoananassae♀* (C) *D.parabipectinata♂ X D. pseudoananassae♀.* (Magnification:40X / Scale bar: 20μm).

### Expression of spermatogenesis genes

11 genes which were previously studied in the *Drosophila* hybrids was considered for the gene expression assay, but 2 were scrutinised and taken out because of uninvolvement of these genes directly into spermatid development or spermatogenesis pathway, we excluded Achintya (achi) and twine because of multiple hits while annotating to design the primers for *D. pseudoananassae*. rest of the 7 genes have shown misxpression in the hybrids of several other *Drosophila* species. Expression studies on *Drosophila bipectinata* were not studied before hence the nature of selecting genes were sceptical and we had to relay mostly on the expression studies of other species hybrids, most of them were found to be on *D. simulans* clade.

We studied testes specific gene expression for the remaining 7 candidate genes selected through gene annotation criteria as discussed above among the 12 hybrid males of *D. bipectinata* species complex. All the candidate genes showed significant misexpression among most of the hybrid species group compared to their respective parents, but reference gene analysed didn’t show expressional difference among the species group from the qRT-PCR data.

Comparing the gene expression data revealed significant level of down regulation in all the hybrid groups relative to the pure species parental groups. Table 1 encapsulates expression results of all the hybrids and pure species through Tukey’s post-hoc test, showing a congruous data nevertheless of the pure species accounted as calibrator, also when Kruskal-Wallis non parametric test was applied to compare the expression of spermatogenesis genes between the hybrids and pure species the results were compatible. For the hybrids of *D. bipectinata* and *D. malerkotliana*, significant underexpression of genes was found among the sterile hybrids except for *cannonball* (*can*) and *mst98ca* (Fig 4) (Table 1). Among the hybrids of *D. bipectinata* X *D. parabipectinata* except *topi* and *can*, rest of the 5 genes didn’t show significant difference in expression compared to parents. Data between the hybrids of *D. malerkotliana* and *D. parabipectinata* indicates a highly significant downregulation of spermatogenesis genes except for can in one of the hybrids in which *D. malerkotliana* served as maternal parent (Fig 5).

**Fig 4.**
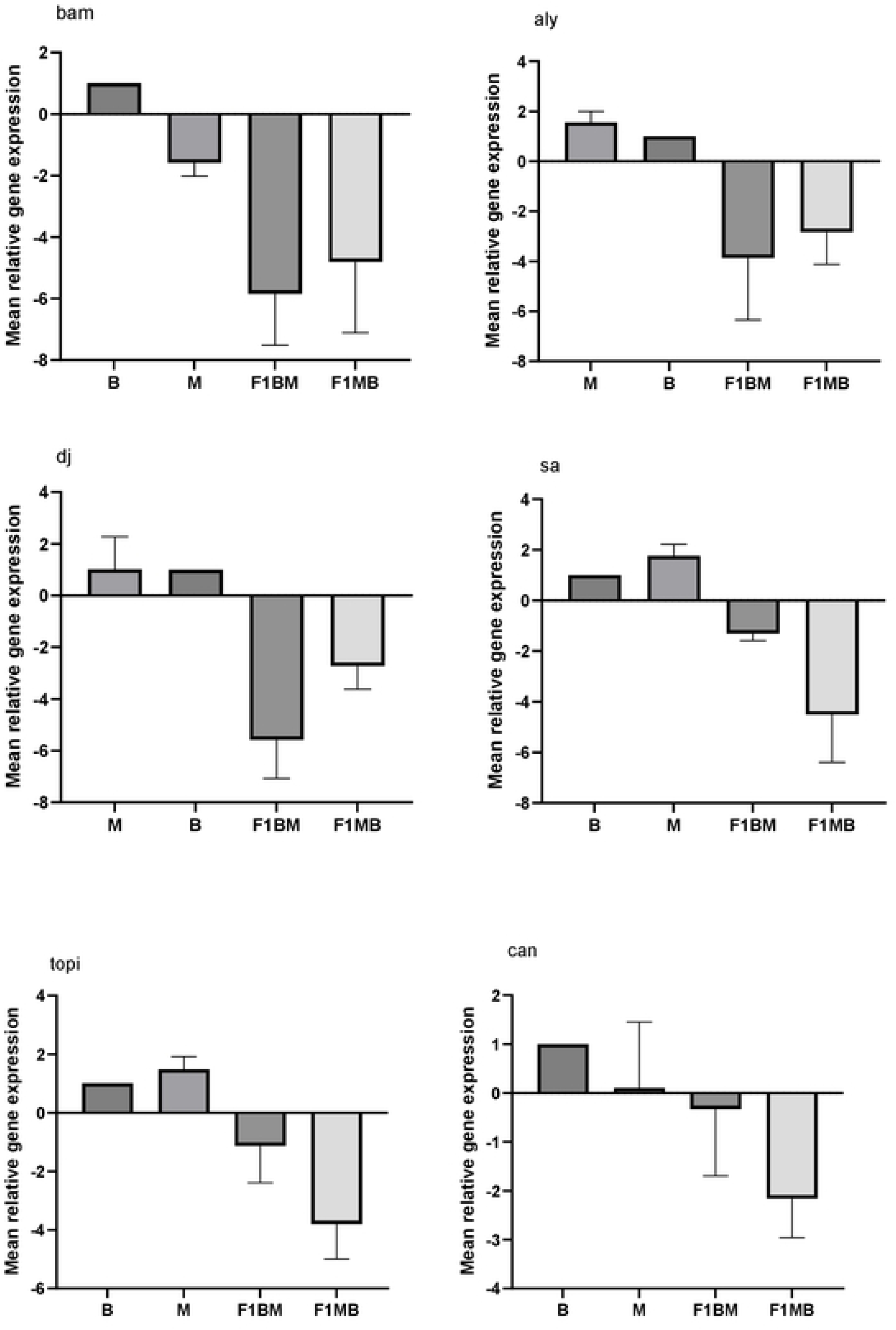
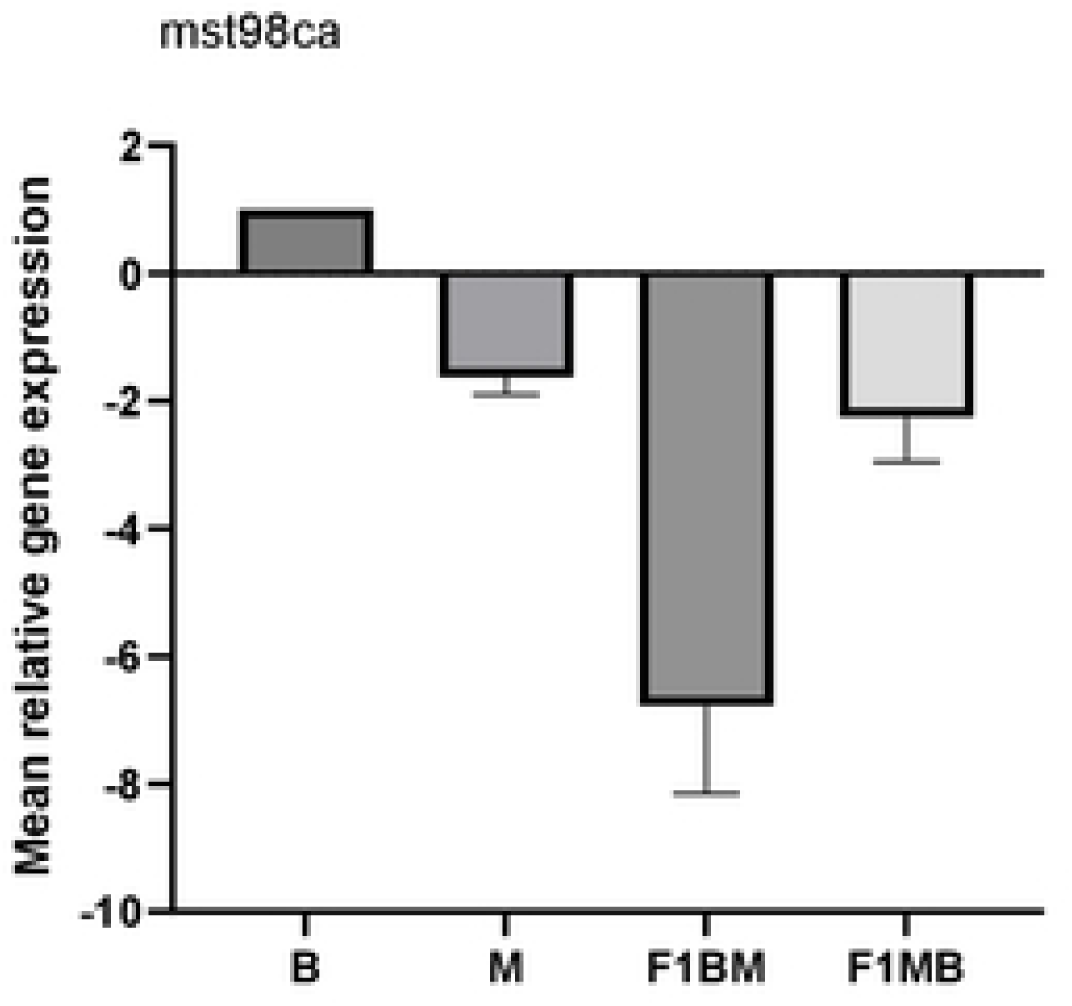
Mean relative gene expression levels of *D. bipectinata* and *D. malerkotliana* (parental pure species) and reciprocal hybrids. Under expression of spermatogenesis genes among the hybrids compared to parents suiting the sterility hypothesis. Error bars represent +_1 standard errors of the mean.

**Fig 5.**
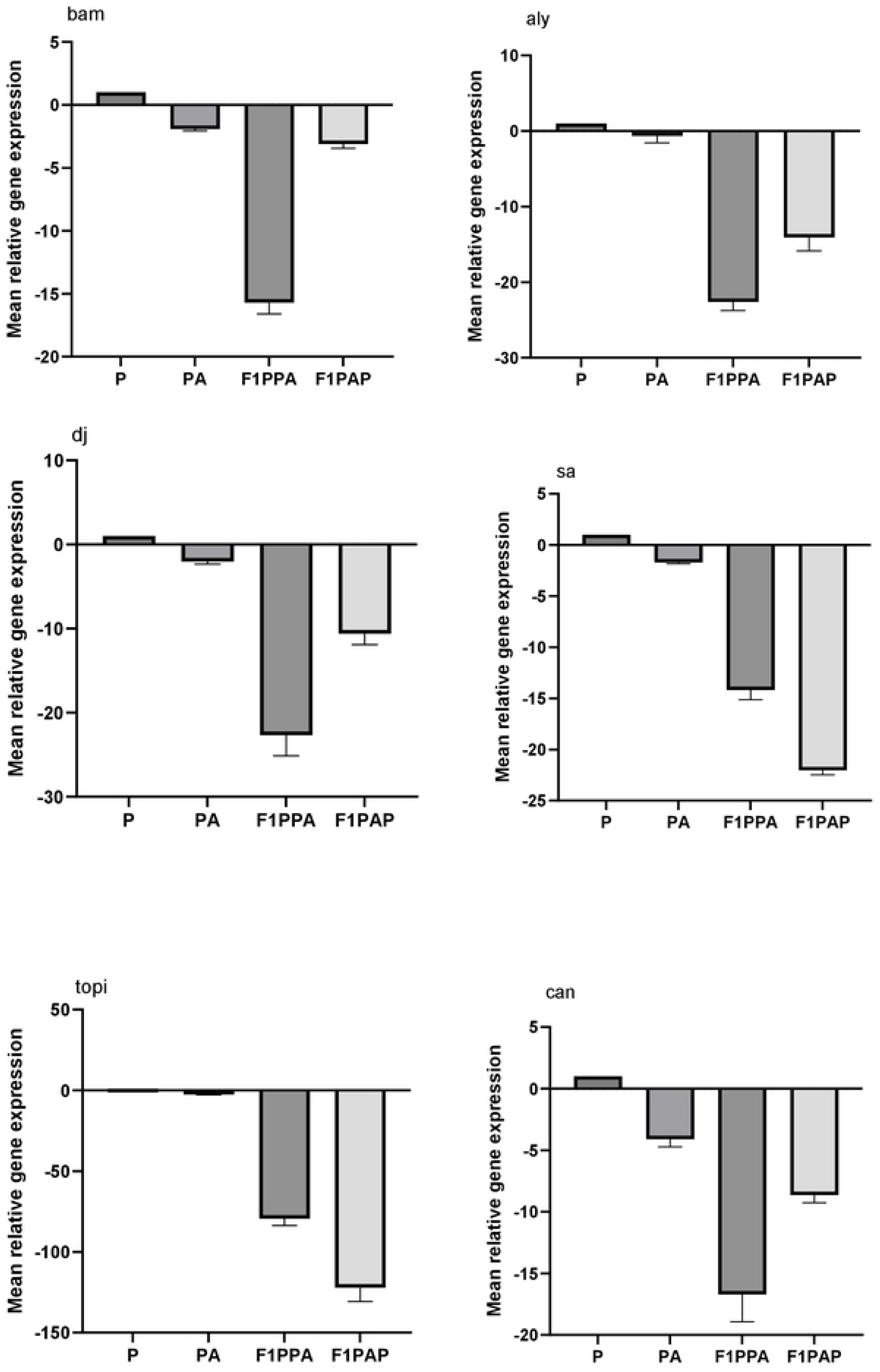
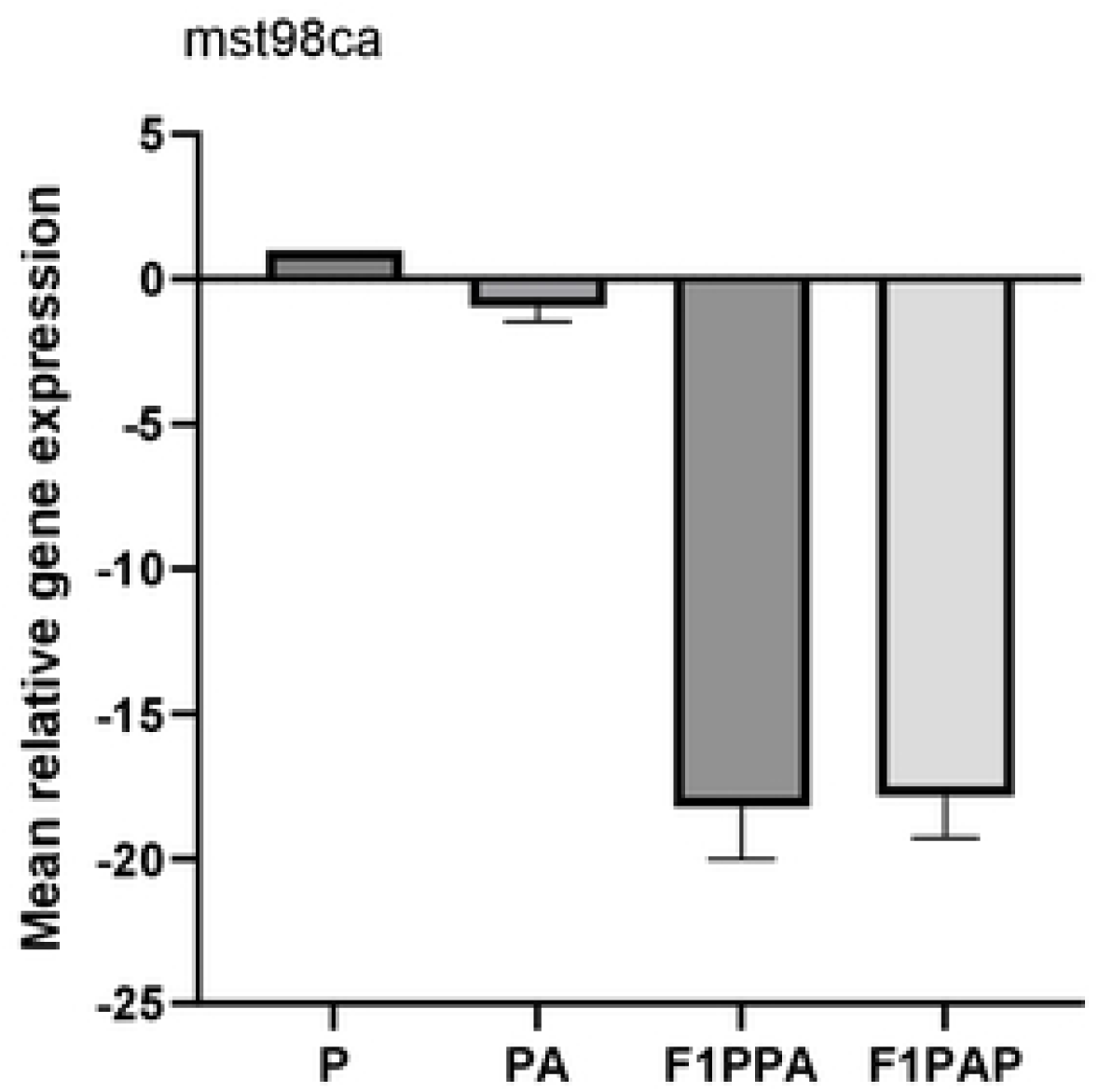
Mean relative gene expression levels of *D. parabipectinata* and *D. pseudoananassae* (parental pure species) and reciprocal hybrids. Under expression of spermatogenesis genes among the hybrids compared to parents, not suiting the sterility hypothesis. Error bars represent +_1 standard errors of the mean.

**Table 1:**
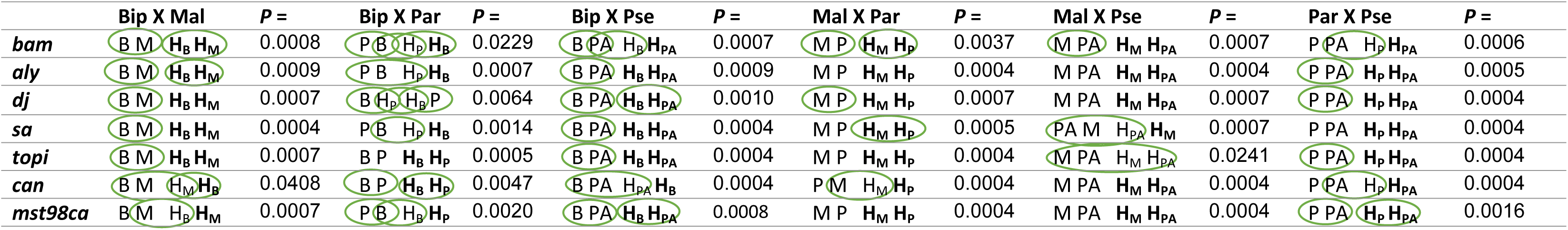
Differences in the expression pattern of spermatogenesis genes among the pure and hybrid species group. Species names are denoted to the first letter of the species name (B = *bipectinata*, M = *malerkotliana*, P = *parabipectinata* and PA = *pseudoananassae*). H represents hybrid and the subscript attached indicates the maternal parent of the respective hybridc. Species groups inside the same circle represents lack of significant differences in the expression of genes (Tukey’s post hoc test). Hybrid species which are denoted in bold letters shows significant differences in expression profile compared to both the parental pure species (Tukey’s post hoc test). FDR-corrected p-values (P) from Kruskal-Wallis test are also represented for comparison (Refer S3 table for abbreviations).

All the hybrids of *D. pseudoananassae* showed significant misexpression excluding few hybrids for *bam*, *sa*, *topi* and *can*. We found no differences in expression of *topi* in the hybrids of *malerkotliana* and *pseudoananassae*, but the rest of the genes showed significant downregulation. We used *RpL32* as control for normalisation and applied one way ANOVA, pair wise comparison test to analyse the expression of genes, the overall results show a significant level of misexpression for all the genes (few exceptions) from the testes samples of both direct and reciprocal hybrids compared to pure parental species. Two germline proliferation genes *bam* and *aly* (Tukey’s post-hoc - P values of *bam* and *aly*: Bip X Mal, P = <0.0001, <0.0001: Bip X Par, P = 0.0045, <0.0001: Bip X Pse, P = <0.0001, <0.0001: Mal X Par, P = <0.0001, <0.0001: Mal X Pse, P = <0.0001, <0.0001: Par X Pse, P = <0.0001, <0.0001) and one spermatoid differentiation gene *Mst98ca* (Tukey’s post-hoc - P values of bam and aly: Bip X Mal, P = <0.0001: Bip X Par, P = <0.0001: Bip X Pse, P = <0.0001: Mal X Par, P = <0.0001: Mal X Pse, P = <0.0001: Par X Pse, P = <0.0001) significantly underexpressed in comparison with their respective parental species groups. One of the meiotic control gene studied here, can, showed significant misexpression in most of the sterile hybrids (Tukey’s post-hoc - P values of *bam* and *aly*: Bip X Mal, P = 0.0016: Bip X Par, P = <0.0001: Bip X Pse, P = <0.0001: Mal X Par, P = <0.0001: Mal X Pse, P = <0.0001: Par X Pse, P = <0.0001), but some of the hybrids shows non significance with their respective parents (Table 1). Hybrids of *D. bipectinata* X *D. malerkotliana*, *D. bipectinata* X *D. pseudoananassae*, *D. malerkotliana* X *D. parabipectinata* shows similarity in the expression pattern with their maternal parent in Tukey’s multiple comparison test. Rest of the genes showed similarly significant lower expression in the hybrids, however some of the sterile hybrids showed non-significantly under expression of genes compared to parents (Hybrids of *D. malerkotliana* X *D. pseudoananassae* for *sa* and *topi*, hybrids of *D. bipectinata* X *D. parabipectinata* for *dj* and *sa*) (Table 1).

Although some of the studies have taken intraspecific fertile hybrids as control to study the spermatogenesis genes misexpression between the hybrids, or even in some of the subfamilies of *Drosophila*, interspecies hybrids produces both fertile and sterile hybrids, corresponding to the hypothesis of gene incompatibility theory, by considering fertile offsprings as the control, *bipectinata* species complex does not produces any of the fertile hybrids or even to consider intraspecies hybrids of *bipectinata*, we don’t have substantial evidence to consider if the spermatogenesis genes of different strains of *bipectinata* complex are evolving rapidly. In our previous study the rate of non-synonymous substitutions was minimal among the different strains of *bipectinata* complex but the substitution rate was only found significant when we compared the variations between different species of the sub-complex but not the strains(Manjunath and Shakunthala, 2023). Hence, we ruled out considering intraspecies fertile hybrids for the study.

### Promoter motif divergence and quantitative binding strength analysis

To explore whether the hybrid misexpression of spermatogenesis genes could be linked to cis-regulatory divergence, we examined the promoter regions of five candidate genes (*aly, bam, dj, sa,* and *topi*) across the four species of the *Drosophila bipectinata* complex. Approximately 1–2 kb of upstream sequence was obtained for each gene, and transcription factor binding sites (TFBS) were identified using FIMO with the JASPAR insect PWM collection.

This analysis revealed a mixture of conserved regulatory inputs and lineage-specific turnover (Figs. 6–7). Several motifs were consistently retained across all four species, such as Hnf4 sites upstream of *sa* and *topi*. However, other motifs exhibited clear species-specific gains and losses. For example, a CG4360 site was present only in *D. pseudoananassae* upstream of *aly*, a pan binding site was unique to *D. bipectinata* in *bam*, and *dj* carried a CG7928 motif also specific to *D. bipectinata*. In *sa* and *topi*, Mad and Blimp-1 motifs were gained in subsets of species while absent in others. The presence/absence matrix across species is summarized in a clustered heatmap (Fig. 6), and linear maps for each gene highlight both conserved elements and lineage-specific turnover (Fig. 7). These findings provide direct evidence that promoters of spermatogenesis regulators are evolving rapidly among closely related species (Supporting Table 4).

**Fig 6:**
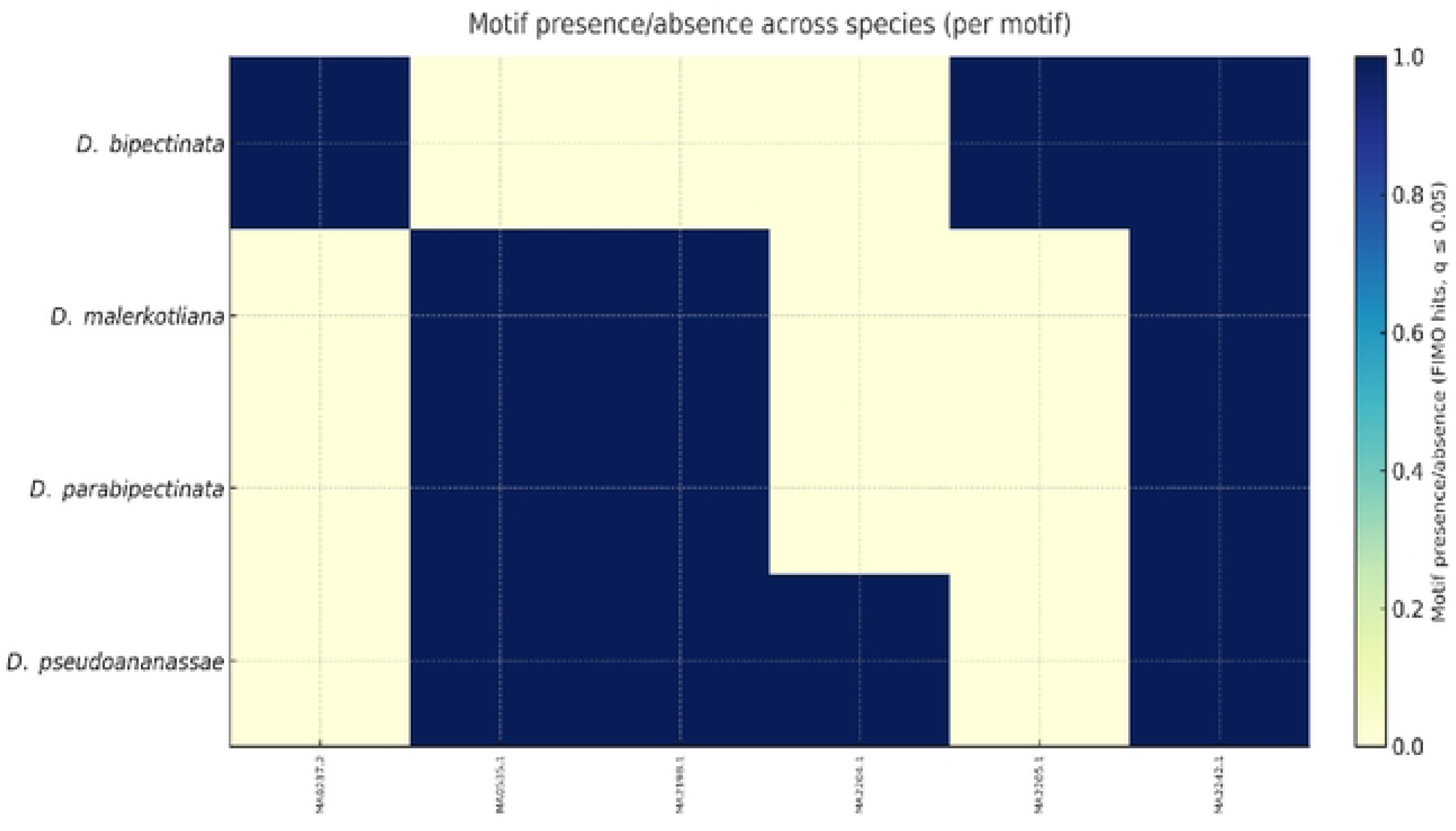
Motif presence/absence heatmap. Heatmap summarizing transcription factor binding site (TFBS) presence and absence across promoter regions of spermatogenesis genes in four species of the *Drosophila bipectinata* complex. Each cell represents detection (q ≤ 0.05, FIMO scan with JASPAR insect PWMs) of a motif upstream of a given gene in a species. Conserved motifs, such as Hnf4, are retained across species, while lineage-specific motifs (e.g., CG4360, pan, Mad, Blimp-1) illustrate cis-regulatory turnover.

**Fig 7:**
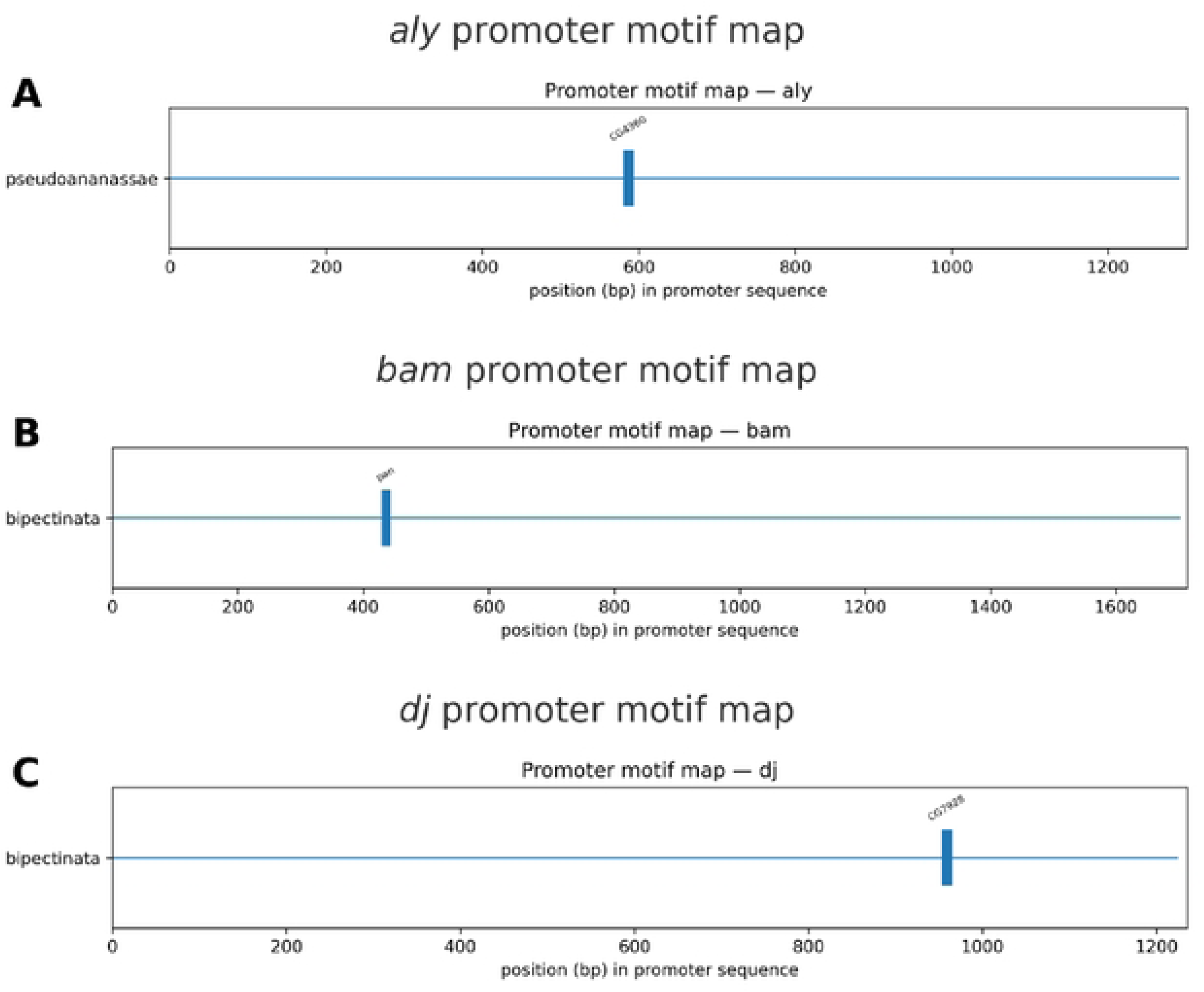
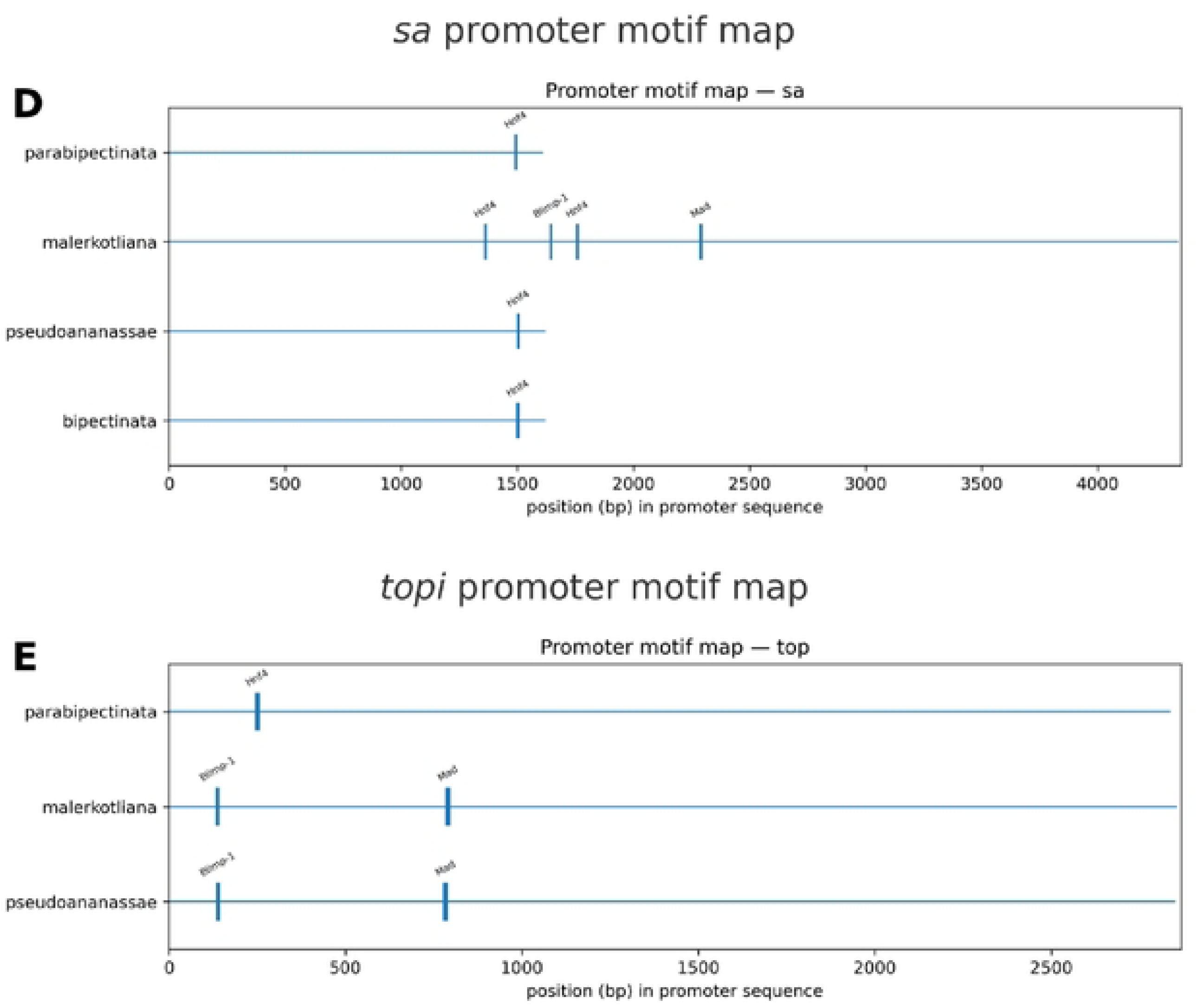
Promoter motif divergence in spermatogenesis genes. Linear motif maps of five spermatogenesis gene promoters (A–E) and the overall motif presence/absence matrix (F) across the *D. bipectinata* complex. (A) *aly*, (B) *bam*, (C) *dj*, (D) *sa*, and (E) *topi* maps show conserved and lineage-specific TFBS detected within ∼1–2 kb upstream regions. For example, a CG4360 site was unique to *D. pseudoananassae* upstream of *aly*, a pan site occurred only in *D. bipectinata* upstream of *bam*, and Mad and Blimp-1 motifs were observed in subsets of *sa* and *topi* promoters.

**Fig 8:**
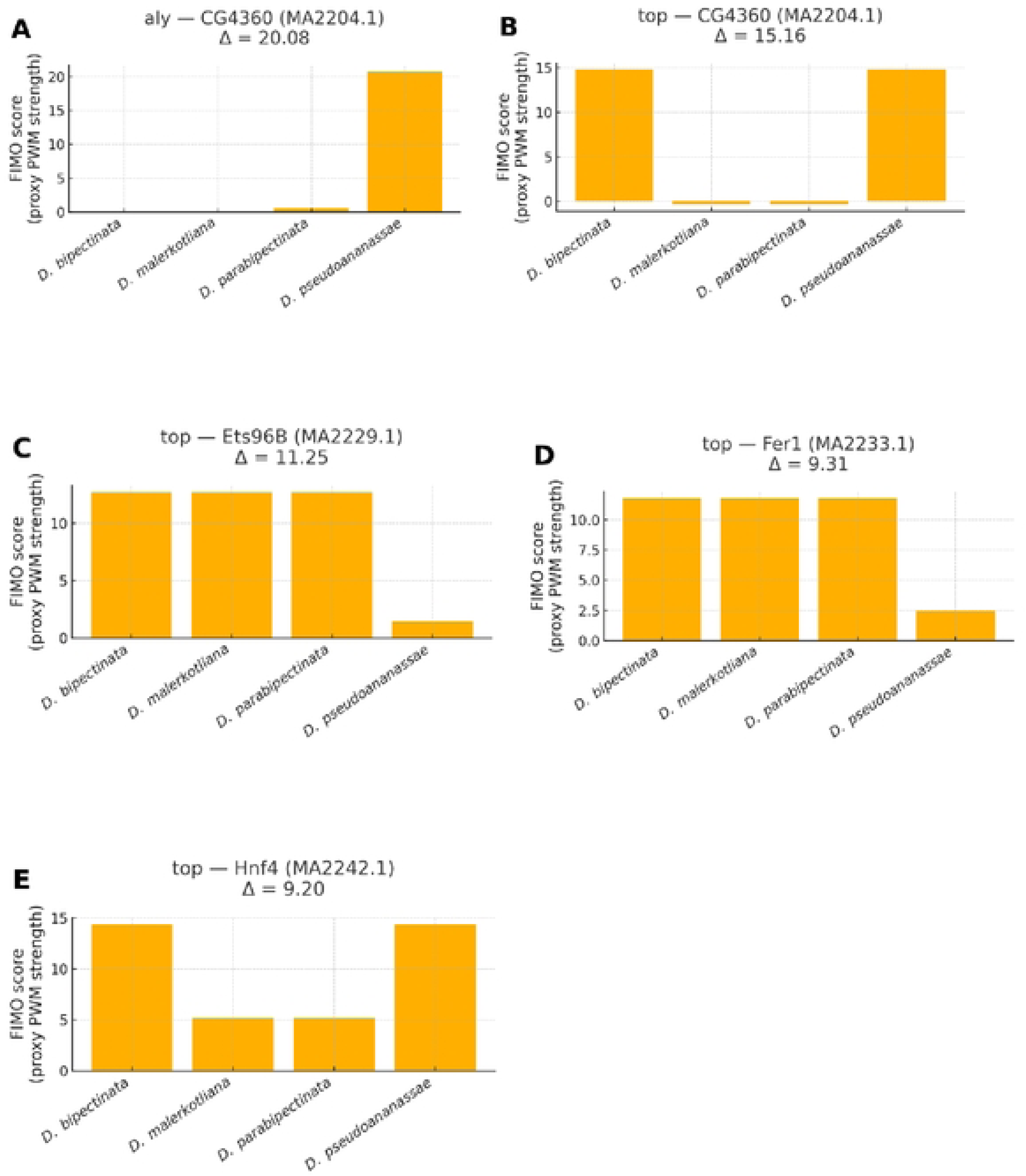
Quantitative divergence in promoter motif strength. PWM-delta analysis of transcription factor binding motifs with the largest interspecific differences in predicted binding strength. Bars show FIMO PWM scores for each species; Δ = max − min score. (A) *aly*—CG4360 motif (Δ ≈ 20.1), (B) *topi*—CG4360 motif (Δ ≈ 15.2), (C) *topi*—Ets96B motif (Δ ≈ 11.2), (D) *topi*—Fer1 motif (Δ ≈ 9.3), (E) *topi*—Hnf4 motif (Δ ≈ 9.2). Large Δ values indicate substantial quantitative cis-regulatory divergence.

Because presence/absence calls alone may underestimate functional divergence, we next examined quantitative differences in predicted binding strength using PWM scores from FIMO hits. For each motif, the highest-scoring site in each species was compared, and the difference between the strongest and weakest score (Δ) was used as a measure of interspecific divergence. This revealed striking quantitative differences in several genes (Fig. 8) (Table 2).

**Table 2:** Quantitative divergence of promoter motifs: Top transcription factor binding sites showing large interspecific differences (Δ > 5) in FIMO PWM match scores. For each gene–motif combination, the species with the strongest and weakest predicted binding strength are shown, together with their scores and the score difference (Δ). High Δ values indicate substantial quantitative cis-regulatory divergence.

| *Gene* | *Motif (TF)* | *Max species* | *Max score* | *Min species* | *Min score* | *$\Delta$ score* |
| --- | --- | --- | --- | --- | --- | --- |
| *aly* | CG4360 | *D. pseudoananassae* | 20.7 | *D. parabipectinata* | 0.62 | 20.08 |
| *bam* | pan | *D. bipectinata* | 18.37 | *D. malerkotliana* | 10.65 | 7.72 |
| *bam* | Ets96B | *D. parabipectinata* | 14.44 | *D. pseudoananassae* | 8.16 | 6.28 |
| *sa* | toy | *D. malerkotliana* | 11.25 | *D. pseudoananassae* | 5.85 | 5.39 |
| *topi* | CG4360 | *D. bipectinata* | 14.82 | *D. malerkotliana* | -0.34 | 15.16 |
| *topi* | Ets96B | *D. bipectinata* | 12.69 | *D. pseudoananassae* | 1.44 | 11.25 |
| *topi* | Fer1 | *D. bipectinata* | 11.75 | *D. pseudoananassae* | 2.44 | 9.31 |
| *topi* | HLH4C | *D. bipectinata* | 14.98 | *D. malerkotliana* | 9.16 | 5.82 |
| *topi* | Hnf4 | *D. bipectinata* | 14.36 | *D. malerkotliana* | 5.16 | 9.2 |

In *aly*, the CG4360 motif was predicted with very high confidence in *D. pseudoananassae* (score 20.7) but was nearly absent in *D. parabipectinata* (score 0.62), producing the largest observed delta (Δ = 20.1; Fig. 8A). *topi* displayed multiple motifs with large interspecific shifts, including CG4360 (Δ = 15.2), Ets96B (Δ = 11.2), Fer1 (Δ = 9.3), and Hnf4 (Δ = 9.2) (Fig. 8B–E). *bam* also showed a strong divergence at a pan motif (Δ = 7.7), while *sa* carried a toy site with a more moderate delta (Δ = 5.4). These analyses demonstrate that cis-regulatory divergence is not limited to motif gain or loss but also involves substantial weakening or strengthening of TFBS across species (Supporting Table 5).

Taken together, the promoter motif turnover and PWM-delta analyses provide direct molecular candidates for cis-regulatory divergence in spermatogenesis genes of the *bipectinata* complex.

The fact that the most divergent motifs occur in genes already known to be consistently underexpressed in sterile hybrids (*bam, aly,* and *topi*) suggests that these cis changes may contribute to the misregulation observed in hybrid testes.

## Discussion

Hybrid male sterility (HMS) is one of the most common forms of postzygotic reproductive isolation (Along with Hybrid inviability, Hybrid female sterility and Hybrid breakdown) in sexually reproducing organisms, which has long been studied as an important barrier to gene flow continuum among closely related species (Coyne and Orr, 2004). In *Drosophila*, lot of studies have deciphered that HMS usually arises from disruption of spermatogenesis dvelopment, through widespread misregulation of germline genes in the hybrid testes (Barbash and Lorigan, 2007; Moehring et al., 2007). While the *Drosophila melanogaster–simulans* clade has been the classical model for studying the genetic basis of HMS for several decades, the *Drosophila bipectinata* species complex has not received much attention despite providing an excellent system for understanding general mechanisms of genetics and evolutionary basis of reproductive isolation. The present study provides the first combined cytological and gene expression analysis of HMS within this complex, and shows that hybrid male sterility is associated with both the developmental arrest of spermatogenesis and with the extensive misexpression of spermatogenesis genes in hybrid males. Along with that, by analysing promoter sequences, we have also identified lineage-specific cis-regulatory divergence, including both motif turnover and quantitative shifts in binding-site strength, which likely contribute to the misexpression of spermatogenesis genes.

Our cytological observations reveals that all the hybrid males suffered from disrupted spermatogenesis, consistent with the complete male sterility observed in heterospecific mating assays. Hybrid males with *D. pseudoananassae* as maternal parent displayed the most severe defects, with testes lacking differentiated spermatids most likely due to early premeiotic arrest, whereas other hybrid combinations progressed further but failed to produce individualized, motile sperm. Such variation in the stage at which spermatogenesis fails has also been reported earlier in the other *Drosophila* clades (Sun et al., 2004; Phadnis and Orr, 2009), and shows that multiple developmental checkpoints are disrupted depending on the genotype shared among the hybrids, reflecting distinct Dobzhansky–Muller incompatibilities acting at different points in gametogenesis.

### The molecular data paralleled with the cytological patterns

Expression profiling of the seven candidate spermatogenesis genes (*bam*, *aly*, *dj*, *sa*, *topi*, *can*, and *Mst98Ca*) showed significant downregulation in most of the male hybrids compared to their parental mRNA expression. Genes such as *bam* and *aly*, which are critical for germline stem cell proliferation and meiotic progression, showed consistent underexpression, giving a mechanistic link to the reduced primary spermatocytes and meiotic arrest observed in the testis of hybrids. Also, Mst98Ca, which is essential for spermatid differentiation, was significantly downregulated, corresponding to the absence of individualized sperm. These results underpin the idea that misregulation of key spermatogenesis pathways crucially contributes to hybrid male sterility. However, not all the spermatogenesis genes followed this pattern; *can* revealed variable expression, and in some of the cases hybrids resembled their maternal parent more closely, consistent with maternal effects and possible nuclear–cytoplasmic incompatibilities (Maheshwari and Barbash, 2011). Such maternal resemblance accentuates the potential importance of regulatory divergence rather than coding-sequence changes alone in driving male sterility.

To investigate whether or not the promoter divergence could underlie these expression changes, we analysed ∼1–2 kb upstream sequences of the five candidate genes across the four parental species. This revealed conserved motifs such as Hnf4 as well as lineage-specific gains and losses, including CG4360 in *aly*, pan in *bam*, CG7928 in *dj*, and Mad and Blimp-1 in *sa* and *topi*. Importantly, quantitative PWM scoring showed that these differences were not only presence/absence changes but also involved large shifts in predicted binding strength. For example, the CG4360 site in aly was strongly predicted in *D. pseudoananassae* but nearly absent in *D. parabipectinata* (Δ score >20), while topi carried multiple motifs with Δ scores of 9–15 across species. These findings indicate that promoter turnover has both qualitative and quantitative dimensions, and provide plausible cis-regulatory candidates for the hybrid misexpression observed (Fig. 3).

A central question which rises by observing these data is whether the current misexpression is a cause of sterility or merely a consequence of disrupted spermatogenesis. Several studies have pointed out this ambiguity, on one hand, consistent with the underexpression of spermatogenesis genes, particularly those expressed post-meiotically, has been documented in many sterile hybrids, suggesting a causal role in spermiogenic failure (Sundararajan and Civetta, 2011; Moehring et al., 2007). On the other hand, fertile hybrids introgressions can also exhibit downregulation of spermatogenesis genes, showing that regulatory divergence alone may generate expression changes even when fertility is unaffected (Ferguson et al., 2013). Thus, although our findings are fully compatible with the misregulation hypothesis, they cannot alone establish causality, which requires genetic mapping, introgression, or allele-specific expression analyses. Evidence from genetic mapping studies underscores this above point. In the *D. mauritiana: D. simulans* system, a small genomic region (HMS1) containing two neighbouring DNA binding protein genes was observed to be contributing quantitatively to sterility, and transgenic rescue with compatible alleles partially restored fertility despite having no consistent differences in transcript abundance (Liénard et al., 2016). This demonstrates that incompatibilities can act through protein function, splicing, or interactions, but not through mRNA levels alone. Applying such mapping and rescue strategies to the *bipectinata* complex would provide direct tests of whether the genes we have identified to be misexpressing are causally involved in sterility.

### Regulatory divergence also plays a central role

Ferguson et al. (2013) showed that autosomal introgressions produced much stronger downregulation of the spermatogenesis genes compared to X-linked ones, insisting that autosomal cis–trans regulatory divergence could be major contributor to misexpression. Our cis-regulatory motif analyses are consistent with this framework: autosomal spermatogenesis genes such as *aly*, *bam*, and *topi* indicate lineage-specific motif turnover and large PWM score deltas, supporting the idea that promoter divergence can generate expression differences independently of coding-sequence change, which implies that some of the expression differences we have observed could result from mismatched regulatory networks in hybrids rather than being the downstream effects of failed spermatogenesis. Allele-specific expression assays in hybrids would allow us to partition cis and trans contributions and clarify the mechanisms. A threshold model has also been proposed in which modest reductions in premeiotic transcript abundance may be tolerated until a certain point, beyond which the deficit causes post-meiotic failure (Michalak and Noor, 2003; Liénard et al., 2016). This framework helps in explaining how the relatively subtle quantitative expression changes can culminate in severe cytological arrest.

Our results also highlight some candidate genes that may represent conserved targets of hybrid dysfunction. The consistent underexpression of *bam* and *aly* in the hybrid males matches the findings with other *Drosophila* clades, where proliferation and meiotic progression genes are often misexpressed in sterile males (Moehring et al., 2007; Sundararajan and Civetta, 2011). The fact that these same genes also exhibit promoter motif divergence and large PWM score differences in our analysis suggests that cis-regulatory changes may underlie a repeatedly disrupted module of spermatogenesis. Previous work has also shown that post-meiotic genes such as Mst98Ca are frequently underexpressed in sterile hybrids (Moehring et al., 2007; Sundararajan and Civetta, 2011), although we did not include this gene in our motif analysis. At the same time, the variability seen for genes like can emphasizes that not all regulatory changes are universal, and that hybrid genotype, maternal effects, and chromosome context can modulate outcomes.

A further perspective comes from comparative work across lineages, for example Gomes and Civetta (2014) showed that misregulation of spermatogenesis genes is often lineage-specific and can be driven either by sterility itself or by rapid male-specific regulatory divergence, consistent with the “faster male” hypothesis. In species pairs with unidirectional sterility, some genes were misregulated only in the sterile hybrid, supporting the sterility hypothesis, while others were misregulated in both fertile and sterile hybrids, suggesting divergence-driven incompatibilities. These studies also points out that different *Drosophila* lineages misexpression are largely seen in non-overlapping sets of spermatogenesis genes, cautioning against the broad generalizations based on the *D. melanogaster* group alone. Our findings of misexpression in the *D. bipectinata* complex, coupled with clear cis-regulatory divergence in promoters, therefore add to the view that although misregulation is a common outcome of hybridization, the specific genes and pathways affected may differ markedly across lineages. This lineage-specificity reinforces the idea that regulatory networks underlying spermatogenesis evolve rapidly, especially for male-biased and X-linked genes, and that the details of hybrid sterility can differ even when the overall phenotype is the same.

Overall, our findings align with other studies including the *D. melanogaster–D. simulans* clade, in which sterile hybrids shows widespread misexpression of spermatogenesis genes (which also produces fertile male hybrids) (Michalak and Noor, 2003; Moehring et al., 2007). The *D. bipectinata* sub-complex, however, differs in that all heterospecific hybrids are sterile, leaving no fertile controls to contrast sterile versus fertile expression profiles. This pattern suggests that incompatibilities have become fixed more rapidly across the complex, potentially reflecting accelerated divergence of spermatogenesis genes or their regulatory networks. Our earlier analyses showing limited nonsynonymous divergence within strains but more substantial divergence between species further support the idea that regulatory divergence, rather than emphasising that protein-coding changes could be underlying the sterility problem. This conclusion echoes the broader view that regulatory divergence often plays a central role in hybrid dysfunction (Landry et al., 2007; Ferguson et al., 2013; Gomes and Civetta, 2014). Our cis-regulatory motif turnover and PWM delta analyses provide direct molecular candidates for such divergence, nominating short promoter elements whose gain/loss, or weakening may drive the expression changes observed among the hybrids. While our qRT-PCR analysis provided targeted evidence of misexpression, focusing on only seven genes which offers a restricted view of the disruption. Broader transcriptomic approaches such as RNA-seq would reveal the full extent of regulatory divergence and could capture small RNAs and chromatin regulators known to disrupt spermatogenesis. Also,Protein-level assays would determine whether transcriptional changes translate into functional deficits, and genetic mapping or transgenic rescue could identify the specific loci responsible. Taken together, these approaches would enable a decisive test of whether the misexpression we observe is cause or consequence of sterility.

A limitation of this study is that we did not directly assess allele-specific expression in hybrids, which would provide unambiguous evidence for the cis-regulatory divergence. Our conclusions are therefore based on indirect evidence from parental promoter divergence and hybrid expression profiles. In addition, two candidate genes (can and Mst98Ca) were not included in our motif analysis due to lack of validated sequences. Although reference sequences exist in databases, we restricted our analyses to experimentally validated Sanger sequences to minimize artifacts. Importantly, the five candidate genes analyzed here, spans distinct functional classes of spermatogenesis regulators, making our observations broadly representative. Future work including hybrid allele-specific assays, expanded promoter analyses, and functional tests of candidate motifs will further clarify the cis-regulatory basis of hybrid sterility. In conclusion, this study demonstrates that hybrid male sterility observed in the *Drosophila bipectinata* species complex is consistently associated with the arrest of key developmental stages of spermatogenesis development which could be due to the misexpression of key spermatogenesis genes. By incorporating the cytological and molecular evidences and the promoter motif turnover and quantitative PWM analysis, we have extended the generalisation of the misregulation hypothesis of HMS problem beyond the *D. melanogaster* - *D. simulans* clade and have highlighted the importance of regulatory divergence, lineage-specific effects, and maternal influences in reproductive isolation. These findings places the *D. bipectinata* species complex within the broader framework of speciation and contribute towards the understanding of molecular mechanisms that restricts in maintaining the species boundaries.

## Statements and declarations

The Author declares that they have no conflict of interest.

## Acknowledgments

We thank the Department of Studies in Zoology and Department of Studies in Genetics and Genomics, University of Mysore for providing the infrastructure. We acknowledge the Council of Scientific and Industrial Research (CSIR), Govt. of India for providing CSIR-NET-JRF/SRF fellowship.

## Supporting information

Supporting table 1: Primers used for PCR amplification and Sanger sequencing the genes

Supporting table 2: Primer sequences used for qRT-PCR

Supporting table 3: Abbreviations and cross setups representing their respective mating types and the hybrids involved in the present study. *H is used for hybrid, with first subscript indicating the paternal parent and the second subscript indicating the maternal parent.

Supporting table 4: Complete list of transcription factor binding site (TFBS) predictions identified by FIMO using JASPAR insect PWMs. Columns include gene, motif ID, transcription factor name, species, binding site coordinates, FIMO score, p-value, q-value, and matched sequence. Only the first 200 rows are shown here; the full dataset is provided in TSV format.

Supporting table 5: Summary of quantitative interspecific differences in predicted binding strength (PWM delta) for all detected motifs. For each gene–motif combination, species-specific maximum and minimum scores are shown along with the calculated Δ. Only the first 200 rows are shown here; the full dataset is provided in TSV/Excel format.

